# Benchmarking overlapping community detection methods for applications in human connectomics

**DOI:** 10.1101/2025.03.19.643839

**Authors:** Annie G. Bryant, Aditi Jha, Sumeet Agarwal, Patrick Cahill, Brandon Lam, Stuart Oldham, Aurina Arnatkevičiūtė, Alex Fornito, Ben D. Fulcher

## Abstract

Brain networks exhibit non-trivial modular organization, with groups of densely connected areas participating in specialized functions. Traditional community detection algorithms assign each node to one module, but this representation cannot capture integrative, multi-functional nodes that span multiple communities. Despite the increasing availability of overlapping community detection algorithms (OCDAs) to capture such integrative nodes, there is no objective procedure for selecting the most appropriate method and its parameters for a given problem. Here we overcome this limitation by introducing a datadriven method for selecting an OCDA and its parameters from performance on a tailored ensemble of generated benchmark networks, assessing 22 unique algorithms and parameter settings. Applied to the human structural connectome, we find that the ‘Order Statistics Local Optimization Method’ (OSLOM) best identifies ground-truth overlapping structure in the benchmark ensemble and yields a seven-network decomposition of the human cortex. These modules are bridged by fifteen overlapping regions that generally sit at the apex of the putative cortical hierarchy—suggesting integrative, higher-order function— with network participation increasing along the cortical hierarchy, a finding not supported using a non-overlapping modular decomposition. This data-driven approach to selecting OCDAs is applicable across domains, opening new avenues to detecting and quantifying informative structures in complex real-world networks.

## 1 Introduction

The complex web of structural connections between brain areas can be represented as a network of nodes (brain regions) and edges (axonal connections) [1, 2]. This representation allows us to quantify and understand properties of the brain’s network structure, like its organization into densely interconnected modules. Connectome modules often correspond to segregated groups of brain regions with distinct, functionally specialized roles [3, 4], wherein highly connected hub nodes are thought to play an integrative role in communicating diverse information between them [5]. Accurately delineating network modules is an essential step in understanding the brain’s modular organization. The community detection algorithms that have most commonly been used to study brain network modules have assumed a hard partition of nodes into mutually exclusive communities; i.e., each node is assigned to a single community. This non-overlapping modular decomposition can be represented as a discrete categorical label that captures each node’s modular affiliation [6].

Applications of community detection to brain networks have ranged from the structural network of individual neurons in *C. elegans* [7] and *Drosophila* [8], to the mesoscopic axonal connections in the mouse [9–11], non-human primate [12], and human [13–15], where inferred network modules align with broad specializations of cognitive function [16]. While these studies support a conceptual model of the brain in which segregated groups of nodes process specialized types of information, hard partitions present limitations that may yield unintended consequences for inferring network-wide modular organization—for example, collapsing two modules bridged by one or more overlapping nodes into a single module. Indeed, transmodal cortical association areas are defined by their flexible multifaceted functions, and might therefore be more accurately affiliated with multiple modules. A representation that allows nodes to belong to multiple communities may thus provide a more complete picture of the brain’s structural and functional organization [17, 18].

Overlapping community detection algorithms (OCDAs) decompose the network into a fixed number of modules, as non-overlapping methods do, but with the additional freedom of allowing each node to belong to multiple communities. A wide range of such methods have been developed [19–24], which may offer a more organic method to characterize the brain’s spatially distributed and fluid network of complex interactions with an overlapping modular architecture. For example, Wu et al. [25] applied the clique percolation method [21] to identify five communities based on gray-matter volume correlations, revealing overlap among well-known functional systems, such as decision-making and emotional processing. Other studies have characterized the hierarchical network architecture of the mouse [26] and macaque [27] structural connectome with algorithms that allow overlapping community assignment, though the overlap itself was an auxiliary point in both studies. By contrast, considerably more work has focused on overlap among functional networks, evaluating functional magnetic resonance imaging (fMRI) [17, 28–35] and/or calcium imaging [18] with a given OCDA to characterize overlapping community structure in functional networks. Collectively, findings from these studies converge on the view that brain areas spanning multiple structural and/or functional modules enable inter-network communication (‘integration’) that still preserves intra-module specialization (‘segregation’)—highlighting how OCDAs offer nuanced information about the rich overlapping patterns in brain community structure that is obfuscated by hard partitions. Moreover, as explored in Bijsterbosch et al. [34], an overlapping node may play a non-stationary role, communicating exclusively with one region at a time or consistently serving as a fully integrative bridging node that facilitates information transfer across functionally specialized systems.

Each of these applications of overlapping community detection to neuroimaging data began with a subjective selection of the OCDA from a range of possible alternatives—without clear evidence of the superiority of one approach over the other for the problem at hand. This speaks to a general challenge in applying community-detection algorithms: each algorithm makes different assumptions about how modules and overlapping nodes (which are members of multiple communities) are defined.

We know that applying an OCDA to a network will always produce *a result*, and that differences in the assumptions between OCDAs yield different assignments of nodes to communities. This multitude of choices for an OCDA algorithm and its parameters thus raises the question: How can we determine which OCDAs may exhibit the most accurate and informative decomposition of a given real-world network?

No single algorithm can perform optimally on all possible data (cf. the ‘no-free-lunch’ theorem [36]); stated in the present context, no community detection method can work well on all networks [37, 38]. Smith et al. [39] developed a ‘Question-Alignment’ approach that tailors the optimal community-detection method based on hypothesized community properties, but this requires *a priori* predictions about community structure without a quantitative benchmark for comparison. Other previous work has benchmarked OCDA performance on typically very large synthetic networks (*>* 1000 nodes) and/or relatively small empirical networks separately [24, 40–45], though algorithmic performance is largely dependent on network topology. As a result, it is difficult to reliably extrapolate previous benchmark comparisons to a network with different characteristics—size, sparsity, topology—such as the brain’s structural connectome. Rather than selecting an OCDA subjectively or from a benchmark study that may not generalize to a new complex network type, we need a way to objectively tailor our choice of algorithm to the structure of a given network at hand.

We address this challenge by developing a flexible method for the data-driven selection of an OCDA for a given empirical network. Here, we focus specifically on the brain’s structural connectome as our target empirical network, with the goal of narrowing down the optimal OCDA and its parameters with which to identify overlapping community structures and interpret their biological significance. Our approach first generates an ensemble of synthetic benchmark networks with properties derived directly from the target empirical network—the diffusion MRI-derived structural connectome from the Human Connectome Project (HCP) [46]—with known ‘ground-truth’ overlapping community assignments, and then compares the performance of a range of OCDAs on these networks. After identifying the top-performing algorithm across the benchmark network ensemble, we then analyze and interpret the resulting overlapping modular decomposition in the context of brain network structure and function. To our knowledge, this is the first work to comprehensively examine overlapping communities in the human brain’s structural connectome using diffusion MRI data. In total, we introduce a novel approach to tailoring OCDA methods to a given real-world network and systematically quantify the overlapping community structure of diverse complex systems.

## 2 Methods

Given an empirical network, our goal is to systematically evaluate candidate OCDAs to identify a top-performing candidate for inferring that network’s overlapping community structure. We propose a datadriven approach with two main steps, as depicted schematically in Fig. 1: (i) *benchmark calibration*; and (ii) *performance evaluation*. First, we generate an ensemble of benchmark networks that mimic community structure properties of the empirical network and contain ground-truth module assignments of nodes. Second, for each generated benchmark network in the ensemble, we quantify how well each OCDA identifies the ground-truth overlapping community structure. This highlights algorithms with strong overall performance on the benchmarks, which are then judged to most accurately uncover overlapping community structure—yielding OCDA(s) tailored to the target network.

**Figure 1:**
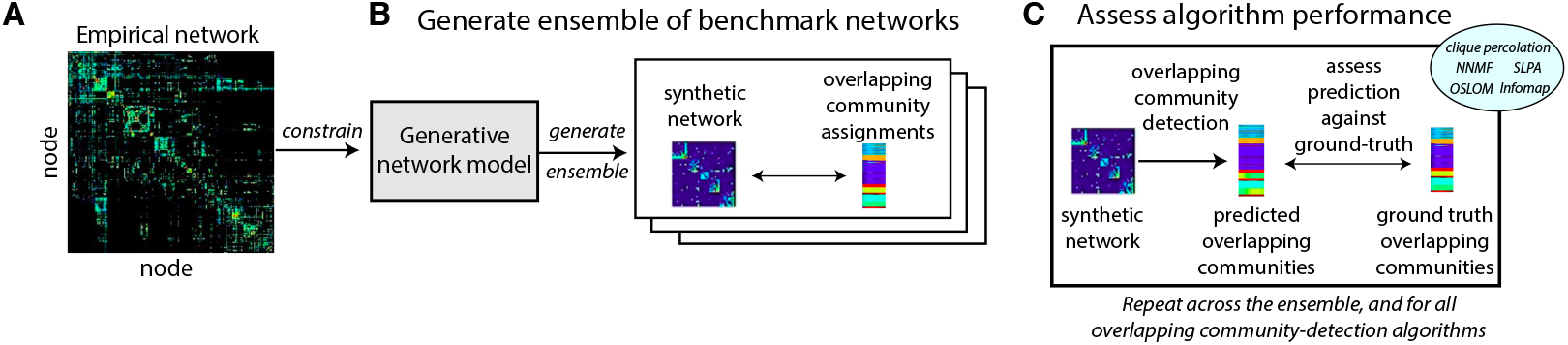
A data-driven method for tailoring overlapping community-detection algorithms (OC-DAs) to empirical networks. **A** An empirical network is shown as an adjacency matrix. **B** We use a generative network model [47] to generate an ensemble of networks with known overlapping community assignments, constrained by the properties of the empirical network. **C** We assess the ability of each OCDA (and parameters) to reproduce the ground-truth overlapping community structure of each synthetic network. Here we compare five OCDAs (clique percolation [21], NNMF [19], SLPA [22], OSLOM [20], and Infomap [48, 49]), across 22 parameter variations, quantitatively evaluating their ability to reproduce the known community assignments using Extended Normalized Mutual Information (ENMI). Algorithms with high ENMI across the ensemble are judged to be most appropriate to infer overlapping community structure in the empirical network.

### 2.1 Generating benchmark networks

The first step in our method involves generating an ensemble of benchmark networks with groundtruth overlapping community assignments, as depicted in Fig. 1B. For the subsequent inference to be informative, these generated benchmark networks should have properties as similar as possible to the empirical network. Here, we focus on weighted, undirected networks (corresponding to our intended application of structural brain networks, such as those estimated using diffusion-weighted imaging [50]). We use the stochastic network generation algorithm of Lancichinetti and Fortunato [47] which—for a given set of parameters that constrain the structural properties of the generated benchmarks—produces weighted networks with overlapping community assignments. Our ability to tune the benchmark networks to match the properties of the target empirical network is controlled by the algorithm parameters, which we describe briefly in following.

Across a given number of nodes, *N*, Lancichinetti and Fortunato [47] benchmark networks are assumed to have a power-law degree distribution, 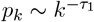, with average degree, ⟨*k*⟩, and maximum degree, *k*_max_. Community sizes are sampled from a power-law distribution with exponent *τ*_2_, with a specific minimum and maximum community count along with a set number of overlapping nodes, *O*_*n*_. Edge weights are assigned under the constraint of a power-law relationship between degree, *k*, and strength, *s*, of network nodes, *s* = *k*^*β*^, determined by the parameter *β*. The extent to which a node’s connections (and their weights) are contained within its own communities (or spread across other communities) is controlled through two mixing parameters: (i) *µ*_*t*_, the average fraction of a node’s edges that are external to the communities it belongs to; and (ii) *µ*_*w*_, the average fraction of a node’s aggregate edge weights external to the communities to which it belongs. Given these parameters, benchmark networks are generated by forming links between nodes within and outside their assigned communities across iterative network rewirings.

To generate benchmark networks with properties that match the empirical network as closely as possible, we require a systematic method to fit the generative parameters of the Lancichinetti and Fortunato [47] algorithm. Some parameters that control the basic connectivity properties of the network (*N*, ⟨*k*⟩ and *k*_max_) were computed directly from the empirical network or—in the case of *τ*_1_ and *β*—were estimated from power-law fitting algorithms [51]. Other parameters governing the community structure (e.g., number and size of communities and overlapping nodes) cannot be inferred from the network but instead require subjective judgment about the type and resolution of community structure that is most appropriate for the application of interest. In the absence of ground truth parameter values grounded in clear domain knowledge (or other priors on the community structure), we sampled these parameters from uniform distributions across reasonable ranges. Specifically, we allowed communities to take any size (from 1–*N* nodes), and sampled the power-law distribution exponent, *τ*_2_ ∼ *U* (2, 4), and the number of overlapping nodes, *O*_*n*_ ∼ *U* (0.1*N*, 0.2*N*), from uniform distributions. To ensure that connectivity was predominantly within communities, mixing parameters were sampled in ranges: *µ*_*t*_ ∼ *U* (0.2, 0.4), and *µ*_*w*_ ∼ *U* (0.2, 0.4). Using this combination of fixed parameters estimated from the network, (*N*, ⟨*k*⟩, *k*_max_, *τ*_1_, *β*), and randomly sampled parameters representing reasonable ranges of the overlapping modular structure of interest, (*τ*_2_, *O*_*n*_, *µ*_*t*_, *µ*_*w*_), we generated an ensemble of 1000 benchmark networks, each resulting from a sampled parameter vector.

The Lancichinetti and Fortunato [47] generative network model makes assumptions about the data that may not hold in all cases; for example, the power-law form of degree distribution that this model assumes will not always be a good representation of real network data. As alternative models are developed in the future that can better fit the key structural properties of a real network, these could be substituted into this benchmark generation step to improve the representativeness of the simulated benchmark networks to the real network data.

### 2.2 Overlapping Community Detection Algorithms

Once we have generated an ensemble of benchmarks that resemble the empirical network (Fig. 1B), we next compare the performance of OCDAs in reproducing the known community structure (Fig. 1C). Here we investigated five different OCDAs, each with various parameter options and delineated briefly below.

*Clique Percolation* [21] generates overlapping communities by clustering a maximal clique network, and then projecting this hard partition of maximal cliques back to an overlapping partition of nodes. A clique is a complete subgraph, and maximal cliques are not subgraphs of other cliques. The method first generates a new network, *G*^*I*^, in which each node is the maximal clique of the original network, *G*, and then computes a hard partition of this new network. As some nodes may be members of multiple cliques, this yields overlapping community assignments in the original network. The algorithm includes a parameter, *k*, that sets the minimum size of maximal cliques to be mapped in the new network, *G*^*I*^. Here we compare seven possible values of this clique size parameter, *k* = 3, 4, …, 9, labeling the resulting algorithms, correspondingly, as: `Clique_3`, …, `Clique_9`.

*Non-Negative Matrix Factorization (NNMF)* [19] uses a Bayesian non-negative matrix factorization model to assign a participation score to each node for each community, which is based on the interaction between that node with other members of the community. NNMF assigns every node a soft membership distribution across communities that can be interpreted as a membership probability of each node to each community. To convert this set of continuous community membership probabilities into a set of discrete labels defining the set of community memberships for a given node, we applied a threshold, *p*_th_. That is, a node is considered a member of a given community if it has a community membership probability in excess of *p*_th_. We compared four thresholds, *p* = 0.1, 0.2, 0.3, 0.4, and labeled the resulting algorithms as: `NNMF_10`, `NNMF_20`, `NNMF_30`, and `NNMF_40`, respectively.

*Order Statistics Local Optimization Method (OSLOM)* [20] is a local method that attempts to find and modify the statistical significance of clusters relative to a global null model in which there is no community structure (the configuration model [52]) by adding/removing neighboring nodes. Thus, OSLOM yields clusters that are statistically unlikely to be found in an equivalent random graph with the same degree sequence; the set of individual clusters found in a network may overlap. In adjusting the significance level, *P*, from which to assess whether to make local changes to a cluster, the resulting size of clusters is changed (low values of *P* yield fewer, larger clusters). We compared a range of ten values *P* = 0.1, 0.2, …, 1.0, denoted as `OSLOM_10`, `OSLOM_20`, …, `OSLOM_100`, respectively.

*Speaker-Listener Label Propagation Algorithm (SLPA)* [22] is an extension of the Label Propagation Algorithm (LPA) to the overlapping setting. In LPA, the community assignments of individual nodes are updated iteratively based on the majority label possessed by its immediate neighbors [53]. In the overlapping generalization, SLPA, nodes are not forced to take a single label, but can hold the memory of previously observed labels; the more frequently a node has observed a node label, the more likely it is to then propagate it to its neighbors. After some maximum number of iterations of label propagation, nodes are assigned to communities based on the most frequently observed labels in their memories. Once communities are identified, SLPA removes labels observed less than the fraction *r*; while *r* values in the range 0.01 to 0.45 have been used previously [54–57], here we set *r* = 0.09 as it sat within the range of thresholds yielding convergent results in the original SLPA publication [22]. Xie et al. [22] note that the set threshold is exclusively for post-processing (i.e., refining communities after they have been defined), such that the SLPA dynamics are determined solely by network topology and employed interaction rules. We denote this method as `SLPA`.

*Infomap* [48, 49, 58] is an information-theoretic method operating under the principle that network communities can be identified by compressing the description of the probability flow of random walks. It solves the map equation, which quantifies the description length of a random walker’s trajectory through the network, by balancing the information required to describe movements between communities and that needed to move within communities. Infomap involves construction of a ‘codebook’, meaning a set of labels (or ‘codes’) assigned to different parts of the network that efficiently capture the trajectories of the random walker within the network. This method was extended to capture over-lapping nodes using a generalized map equation that allows these codebooks to allow multiple modules assigned toa single node [58]. We denote this algorithm as `Infomap`.

In our benchmark comparison procedure, outlined in the following sections, the five OCDA methods and corresponding parameter combinations yielded a total of 22 specific OCDA algorithms to evaluate.

### 2.3 Performance evaluation

After applying each OCDA to predict overlapping community structure across nodes within each benchmark network, we next assess how well each algorithm reproduces the ground-truth overlapping community assignment, as depicted in Fig. 1C. We quantified performance using the Extended Normalized Mutual Information (ENMI) [59], which measures the overall similarity of two overlapping community assignments across all nodes. ENMI is an extension of normalized mutual information— which was shown to be important for community detection in finite sample sizes in Newman et al. [60]—that applies to overlapping modular assignments. A larger ENMI value for a given algorithm (and particular parameter value, when applicable) indicates superior recovery of the underlying overlapping community structure in the benchmark network. The best algorithm judged by these metrics is then applied to the real-world network of interest to obtain its underlying overlapping community structure.

### 2.4 Preprocessing structural connectome data

We apply the systematic OCDA selection method to structural connectome data collected from 973 participants from the Human Connectome Project (HCP) [46]—comprising a subset with complete diffusion MRI data from the broader S1200 release [61]. All participants were healthy and aged between 22 and 35 years and provided written informed consent; ethics was approved by the Institutional Review Board of Washington University in St Louis [46]. Structural magnetic resonance imaging (MRI) data was acquired using a 3T Siemens Skyra scanner with a customized head coil (100 mT/m maximum gradient strength and a 32-channel head coil) located at Washington University in St. Louis, Missouri. Diffusion-weighted images were acquired using a spin-echo echo planar imaging sequence and processed according to the HCP Diffusion pipeline [62], with additional preprocessing using `MRtrix3` [63, 64] and the `FMRIB` Software Library [65]. Tractography was conducted in each participant’s structural MRI (T1) space using second-order integration over fibre orientation distributions (iFOD2), a probabilistic algorithm that improves the quality of tract reconstruction in both highly curved and crossing fibre regions [66]. Anatomically Constrained Tractography (ACT) and Spherically Informed Filtering of Tractograms (SIFT-2) were applied to the tractography data [67] to further improve the biological accuracy of these structural networks. Here, we focused on the cortical right hemisphere using the 180-region HCP-MMP1 parcellation from Glasser et al. [68]. A representative group-level connectome was calculated from the individual-level data using consistency-based thresholding [69], retaining only the weights from edges that ranked in the lowest 15% for weight coefficient of variation (CV) across the entire cohort. This method retains the most stable edge weights across the entire sample, with edges failing to meet this cutoff threshold set to zero. Of note, one node—*13l*, in the orbitofrontal cortex—exhibited highly variable edge weights (all with CV of 0.3 or higher), and had a total degree of zero (after applying the 0.15 threshold-based consensus). To prioritize consistency in edge weights for community detection, we thus excluded region *13l* from our analysis. In this group-level connectome, edge weights correspond to the thresholded average number of streamlines between the two regions terminating within a 5 mm radius of each other. Due to the range in streamline count magnitudes across edges, that spanned multiple orders of magnitude, we log-transformed the right-hemisphere edge weights between region–region pairs before further analysis.

### 2.5 Applying OSLOM to the empirical structural human connectome

Since OSLOM was identified as the top-performing OCDA in the benchmark ensemble analysis, we further refined its threshold parameter (*P*) for the applied case study analysis with the empirical right-hemisphere cortical connectome. We compared modular decomposition at each *P* = 0.01, 0.02, … 1 to identify a suitable range of *P* values, across which OSLOM identified a consistent number of modules and overlapping nodes. Importantly, OSLOM is a stochastic algorithm, such that the resulting decomposition can vary depending on the user-supplied seed for the pseudo-random number generator (which determines which nodes are initially selected as the basis for community evaluation) [20]. Most prior applications of OSLOM to interdisciplinary real-world problems that specify *P* and algorithm hyperparameters do not specify the random seed used to generate their results (in which case the seed is automatically generated based on the time at analysis, down to the microsecond) [70, 71]. Bruner et al. [72] used the singular pre-specified seed of 73, and Acman et al. [73] examined a handful of random seeds (1, 5, 42, 93, 212) before manually selecting results obtained with an individual seed (42). Gates et al. [74] did not specify whether seeds were set, but they applied OSLOM to the functional brain connectome 10 times per participant, deriving an ‘element-centric similarity matrix’ based on similarity of individual OSLOM decompositions between participants.

Recognizing that the robustness of a particular connectome decomposition is integral for inferring generalizable biological significance in the human connectome, we systematically compared across 100 different seeds (specifically, setting the seed to each of the integers from 1, 2, … 100) for each *P*-value threshold. We confirmed that repeating OSLOM partitioning with the same seed and threshold (*P*) yielded the same outputs, mitigating the stochasticity issue. Once the *P*-value range was narrowed down to *P* = 0.15, 0.16, 0.35 based on the mean number of modules and overlapping nodes (averaged across the 100 seeds per threshold, cf. Figs S1A-B), we compared the ENMI between each seed– seed pair per *P*-value threshold (cf. Fig. S1C). For each of the 100 seeds, we computed the mean ENMI between that seed and all other seeds, identifying 61 as the seed with the highest overall ENMI—meaning it was maximally representative of decompositions resulting from all other seeds. After selecting a final value of *P* = 0.3, as an intermediate value that yielded top performance in the simulated benchmark analysis, we confirmed that seed 61 is highly representative of all 99 other evaluated seeds at *P* = 0.3, with a mean seed–seed ENMI of 0.64 ± 0.11 (compared to the average across all other seeds of 0.59 ± 0.1). All subsequent analyses in the empirical cortical connectome case study pertain to the modular decomposition obtained from *P* = 0.3 with seed 61. Taken together, while stochasticity in community detection algorithms is challenging, we developed and implemented a systematic approach from which we selected a representative result from OSLOM to subsequently analyze.

### 2.6 Comparing overlapping nodes with biologically relevant properties

In order to investigate various aspects of how the overlapping community structure relates to the putative functional hierarchy in the cortex, we used principal gradients (PGs) of functional connectivity [75]. We obtained pre-processed maps of the first and second PGs (PG1 and PG2, respectively) from the *neuromaps* python package [76] (version 0.0.5). After transforming the reference maps into `fsaverage` surface space, we analyzed the vertex-wise values for PG1 and PG2 for each of the 180 regions in the HCP–MMP1 parcellation [68]. We summarized the mean values across vertices to obtain region-averaged values for PG1 and PG2, respectively.

### 2.7 Comparing overlapping and non-overlapping modular partitions

Non-overlapping community detection methods, like the Louvain method [77], yield modular network decompositions from which nodal statistics that capture aspects of connection diversity across the modules, such as the participation coefficient [78] and participation entropy [79, 80]. To clarify whether the top-performing OCDA can provide additional and unique information beyond that of a non-overlapping partition, we compared network properties from the two modular decomposition types. For a representative non-overlapping partition, we applied Louvain community detection using the method from Rubinov and Sporns [81], first tuning the resolution parameter *γ* to yield relatively consistent decompositions across 100 iterations, each with the seed set to an integer from 1, 2, … 100 (as with OSLOM). After sweeping across *γ* values ranging from 0.5 ≤ *γ* ≤ 1.5, we selected *γ* = 1 (which happens to be the default in the Brain Connectivity Toolbox [81]) as this forms an elbow in the number of communities with lower variance across seeds than higher *γ* values (as plotted in Fig. S5A). We selected a representative seed (98) based on mean normalized mutual information (NMI) with all other seeds (total of 100, as shown in a heatmap in Fig. S5B), which yielded a final six-community decomposition—projected onto the right hemisphere cortical surface in Fig. S5C. As a robustness test, we also compared results using *γ* = 1.5 to enforce a seven-module (non-overlapping) decomposition for the direct comparison of seven communities between OSLOM and Louvain, with the results shown in Fig. S5D-G.

We computed two frequently-used nodal measures from the Louvain decomposition using the python implementation of the Brain Connectivity Toolbox [81] (`bctpy`, https://github.com/aestrivex/bctpy): (i) within-module strength (*z*-scored), *z*, which measures how strongly a node is connected to others within its own community; and (ii) participation coefficient, *P*, which measures the diversity of a node’s connections across network modules [2, 78, 80]. To understand the relative position of each OSLOM-defined overlapping node within the non-overlapping Louvain partition, we plotted all nodes in *z* − *P* space, following the characterization in Guimerà and Nunes Amaral [78]. For direct comparison with the top-performing OCDA, we adapted the algorithm for *P* to allow each overlapping node to be assigned to multiple modules.

### 2.8 Data visualization and code availability

The overlapping community decomposition was visualized within the parcellated cortical surface using the *ggseg* package in R [82]. We used the *NetworkX* package [83] in Python to visualize community structure with a ‘circos’ layout and an undirected graph network layout in Fig. 4B. Raincloud plots are visualized using the ‘see’ [84] and ‘ggplot2’ [85] packages in R. An open and extendable Matlab implementation of our methodology is available on GitHub^1^, as is code to reproduce our main results in a combination of Matlab and Python (and R, for visualization purposes only)^2^.

## 3 Results

Here, we apply our data-driven algorithm selection method to characterize the overlapping modular structure of the right cortical connectome, derived from 973 individuals in the HCP dataset [46]. As depicted schematically in Fig. 1, our systematic approach generates an ensemble of benchmark networks with properties that mirror the target empirical connectome, with a ground-truth overlapping structure that enables selection of the top-performing OCDA. After identifying OSLOM as the top-performing algorithm across the benchmark ensemble, we then analyze the overlapping community structure inferred by this method, comparing the resulting decomposition with functional properties to demonstrate the unique insights beyond those that can be inferred from a traditional non-overlapping partition.

### 3.1 The connectome-tailored benchmark network ensemble highlights OSLOM as the top-performing algorithm

For the 180-node right-hemisphere human structural connectome, shown in Fig. 2A, we followed the general procedure depicted in Fig. 1. We first needed to set the benchmark-generation parameters, (*N*, ⟨*k*⟩, *k*_max_, *τ*_1_, *τ*_2_, *β, O*_*n*_, *µ*_*t*_, *µ*_*w*_), to match the data, as described in Methods (Sec. 2.1). Basic network parameters were computed directly from the network: *N* = 180, ⟨*k*⟩ = 29, and *k*_max_ = 102. Plotting node degree, *k*, against node strength, *s*, shown in Fig. 2B, revealed *s* ∝ *k*, allowing us to identify the exponent in *s* ∝ *k*^*β*^, as *β* = 1. A limitation of the Lancichinetti and Fortunato [47] algorithm is its assumption of a power-law degree distribution, 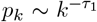 which, as shown in Fig. 2C, does not hold for this structural connectome. We nevertheless set *τ* as the best power-law fit to this distribution, as *τ*_1_ = 2. Sampling from all other parameters across appropriate ranges (using uniform distributions, as described in Sec. 2.1), we generated an ensemble of 1000 structural connectome benchmark networks, each with a ground-truth overlapping community assignment.

**Figure 2:**
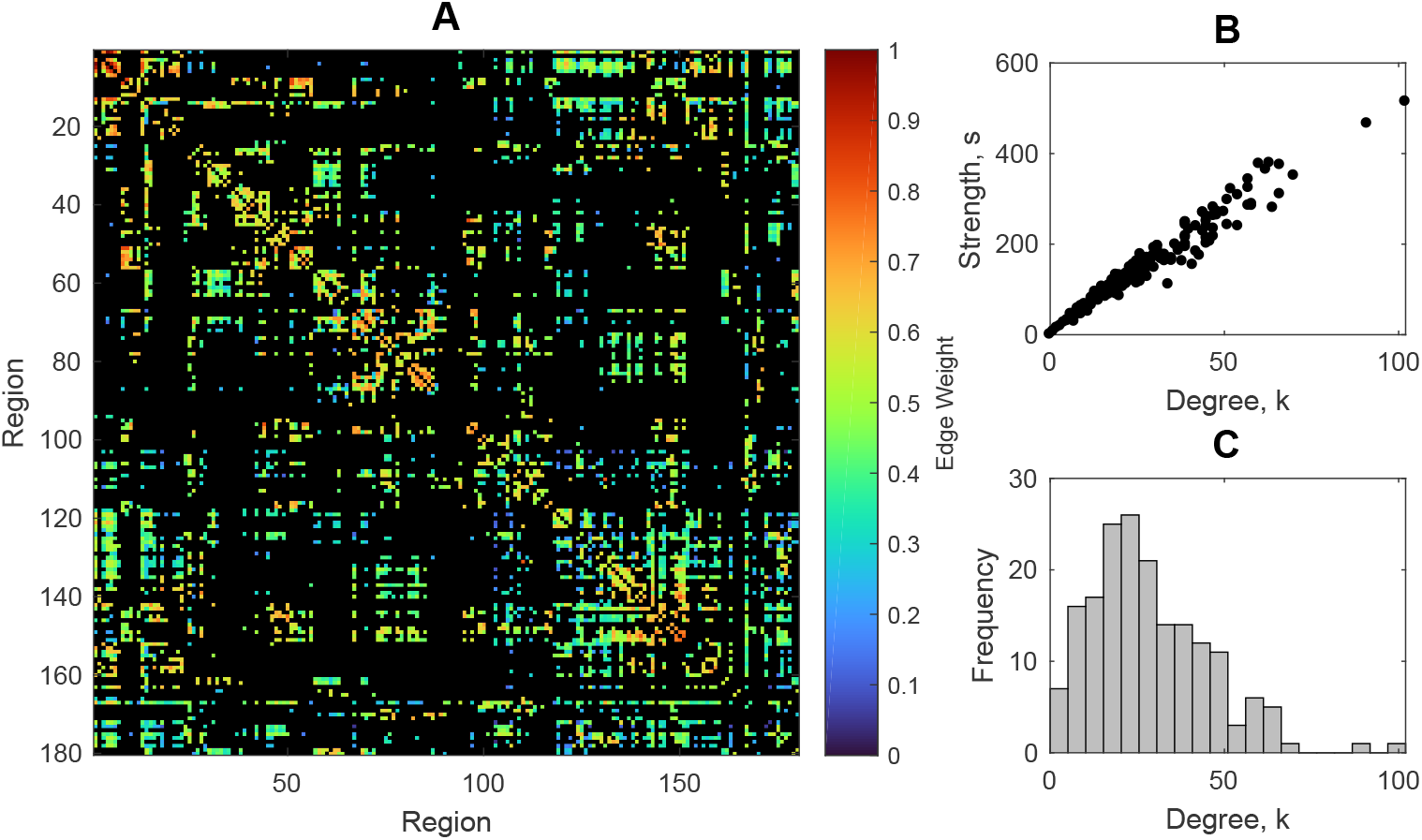
Network properties of the group-level structural connectome of the right hemisphere of the human cortex. **A** The structural connectome of the right cortical hemisphere is shown as a weighted 180 × 180 adjacency matrix, with color representing the log-transformed number of connections (edges) between nodes (normalized to the unit interval for visualization). **B** Plotting node strength (*s*, the sum of weighted connections to the given node) vs. node degree (*k*, the sum of binarized connections) demonstrates that *s* ∝ *k*—enabling inference that *s* ∝ *k*^*β*^, with *β* = 1, for Lancichinetti and Fortunato [47] benchmark network generation. **C** The frequency distribution for degree (*k*) values across nodes, which does *not* exhibit power-law scaling.

We evaluated the performance of 22 OCDAs (based on five representative algorithms) on each generated benchmark network using the ENMI metric (cf. Sec. 2.3). To depict this method, in Fig. 3A we show one representative benchmark network from the 1000 generated with the Lancichinetti and Fortunato [47] method. The ‘ground-truth’ community structure and overlapping community assignments are shown in Fig. 3B, along with those predicted by each of the 22 OCDAs. Visually, we see that variations of the OSLOM method best recapitulate the known overlapping community structure of this representative benchmark, though Clique Percolation results also exhibit similar structure to the ground-truth decomposition. To summarize across all 1000 generated benchmark networks, in Fig. 3C we show the ENMI distributions as raincloud plots for each OCDA configuration (excluding OSLOM with parameter settings that yielded redundant decompositions). The three OSLOM algorithms yielded the highest ENMI values on average, indicating that OSLOM best uncovered the community structure across human structural connectome-like benchmark networks out of the methods compared here.

**Figure 3:**
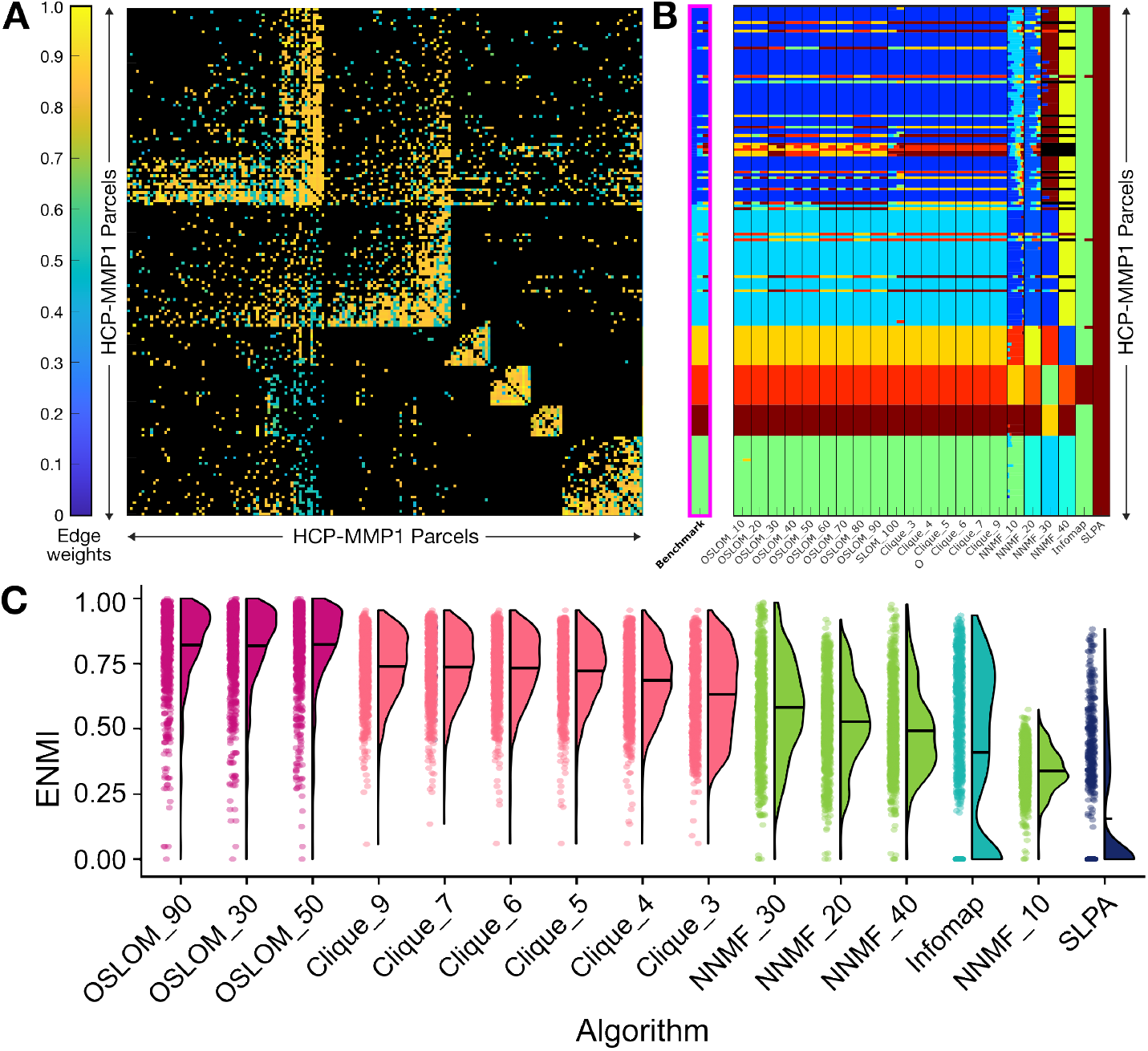
Overlapping community structure predicted by OCDAs (across 22 different combinations of parameters for five types of OCDA methods) on a representative connectome-like benchmark network. **A** The 180 × 180 group structural connectivity adjacency matrix, with the HCP-MMP1 regions [68] sorted according to the ‘Benchmark’ community assignment. Matrix cells are colored according to the consensus-thresholded average edge weights, corresponding to log-transformed streamline normalized to the unit interval [0, 1] for visualization. **B** The ‘Benchmark’ column (highlighted in pink with bold text) represents the ground-truth community structure, in which each of the 180 nodes is assigned to one or more colors. Nodes belonging to the same community are assigned the same color, and overlapping nodes are assigned multiple colors. The other 22 columns on the right depict predictions of the overlapping community structure from each OCDA. Note that colors designate different communities within each OCDA column, but individual colors are not directly comparable between OCDA results (i.e., between columns). **C** For each algorithm, ENMI performance distributions are shown as raincloud plots across the generated ensemble of 1000 benchmark networks. Algorithms are sorted by mean ENMI across the ensemble (shown as black horizontal lines), and algorithms are color-coded as OSLOM (magenta), Clique Percolation (light pink), NNMF (green), Infomap (teal), and SLPA (navy blue). Note that OSLOM results are shown here for three selected resolutions (*P* = 0.3, 0.5, 0.9); ENMI distributions are similar for other values of *P*.

Clique Percolation algorithms exhibited the second-highest mean ENMI values, with performance saturating for clique sizes *k >* 4. Within NNMF, maximal ENMI values were achieved with probability thresholds (*p*) *>* 0.1, while Infomap and SLPA exhibited generally weaker performance. Given these results, we progressed with OSLOM as the preferred overlapping community-detection algorithm to study the structural architecture of this cortical connectome dataset.

### 3.2 Overlapping community detection reveals regions that bridge multiple networks and sit high in the cortical hierarchy

Having identified OSLOM as the top-performing algorithm for uncovering the overlapping community structure in our structural connectome dataset, we progressed with this algorithm to characterize its overlapping community structure. As described in Sec. 2.5, we performed a more comprehensive parameter sweep (across *P*-value thresholds in finer increments of 0.01 across 100 initialization sweeps), and identified a plateau in the number of modules detected (6 to 7) and the number of overlapping nodes detected (14 to 15) between 0.15 ≤ *P* ≤ 0.35 (cf. Figs S1A,B). We selected an intermediate value within the stable plateau of *P* = 0.3, which also yielded among the highest ENMI with the groundtruth overlapping community structure in the benchmark analysis (cf. Fig 3C). After examining pairwise ENMI value across initialization seeds to identify an optimally representative decomposition, we selected seed 61 (highlighted in the ENMI heatmap in Fig. S1), which exhibited a mean ENMI of 0.64 ± 0.11 with all other examined seeds at *P* = 0.3. This selection of parameters is referred to hereafter as `OSLOM_30`.

As shown on the brain’s surface in Fig. 4A, `OSLOM_30` yielded an overlapping modular decomposition with seven modules, which we labeled as follows: (1) dorsomedial cortex; (2) limbic; (3) visual; (4) ventral attention and visual stream; (5) somatomotor; (6) insula; and (7) frontal pole. Of note, most individual regions within each structural community are spatial neighbors, with modules thus forming spatially contiguous clusters. This is consistent with prior findings that spatially proximal regions are more likely to exhibit axonal connectivity between each other [86] and share similar connectivity patterns with the rest of the brain [87, 88]. The seven `OSLOM_30`-defined communities show substantial overlap with the seven canonical resting-state functional networks described in [89] (see Fig. S3 for overlap proportions shown as a heatmap), which is consistent with work from Chen et al. [90] showing that anatomical community structure, based on cortical thickness (not structural connectivity), is closely related to functional systems [90].

Relative to non-overlapping methods, a key benefit of OCDAs is their ability to identify overlapping nodes—i.e., brain regions that bridge two or more different communities. Our `OSLOM_30` decomposition identified fifteen overlapping brain regions: *PreS, ProS, PeEc, PHA1, POS2, RSC, TE1a, PH, STSva, PFm, PGs, 8Av, 8BM*, and *9m*. These overlapping regions are highlighted in bold on the cortical surface along network edges in Fig. 4A and the communities they span are summarized in Table 1. Specifically, these overlapping regions were collectively assigned to six pairs of communities (e.g., ‘Frontal pole + limbic’, ‘Somatomotor + ventral attention and visual stream’), and one region was assigned to three communities (‘Limbic + visual + dorsomedial’). The connectogram (with logtransformed streamline counts, cf. Sec. 2.4) in Fig. 4B verifies that these overlapping regions exhibit structural connections across multiple modules. Examining structural connectivity specifically between these fifteen overlapping nodes in Fig. 4C reveals strong interconnectivity, consistent with the notion that overlapping regions integrate information across segregated brain networks—noting that these regions exhibit higher overall degree on average (cf. Fig S2A).

**Table 1:** The fifteen nodes that OSLOM_30 identified as overlapping across two or more structural communities in the cortical connectome. The ‘+’ sign denotes an overlap between the previous and subsequent listed communities.

| Brain Region | Communities Bridged |
| --- | --- |
| PreS | Visual + limbic |
| ProS | Visual + limbic |
| PeEc | Visual + limbic |
| PHA1 | Visual + limbic |
| POS2 | Visual + limbic |
| RSC | Dorsomedial + visual + limbic |
| TE1a | Visual + ventral attention and visual stream |
| PH | Visual + ventral attention and visual stream |
| PGp | Visual + ventral attention and visual stream |
| STSva | Visual + ventral attention and visual stream |
| PFm | Somatomotor + ventral attention and visual stream |
| PGs | Somatomotor + ventral attention and visual stream |
| 8Av | Somatomotor + frontal pole |
| 8BM | Frontal pole + dorsomedial |
| 9m | Frontal pole + limbic |

Figure 4D plots the overlapping regions on the cortical surface, revealing that the fifteen nodes are evenly distributed along both anterior–posterior and medial–lateral axes. On average, each overlapping node comprised a larger surface area than a non-overlapping node (unpaired Wilcox rank-sum test, *P* = 0.03; cf. Fig. S2B). Comparing the locations of these overlapping regions with those of the seven structural communities, we find that they sit at the spatial interface between adjacent communities— consistent with similar findings reported in the macaque structural connectome [27]. For example, the ‘Somatomotor + ventral attention’ overlapping regions, *PFm* and *PGs*, sit at the ventral end of the ‘Somatomotor’ module and the dorsal end of the ‘Ventral attention and visual stream’ module. Taken together with the inter-module connectivity depicted in Figs 4A,B, this suggests that overlapping nodes can form a structural bridge for rapid integrative processing between the constituent brain networks—in which case these regions could serve as cross-network ‘integrators’ or ‘hubs’ [91–93].

Based on the seminal work in Mesulam [94]—and mounting evidence that ‘integrative hubs’ contribute to multiple functional domains [17, 27, 95, 96], we hypothesized that overlapping regions that bridge two or more structural communities would sit higher in a topographical hierarchy of the cortex, such that their cross-community connectivity enables the exchange of information from distributed modules. To test this hypothesis, we first examined the extent to which each overlapping node mapped to each of the seven canonical resting-state functional networks from Yeo et al. [89]. As shown in Fig. 5A, we found that roughly half of the overlapping nodes map primarily to the higher-order default mode and frontoparietal (cognitive control) systems, which comprise association and paralimbic cortices. By contrast, the remaining overlapping regions correspond primarily to the visual network (a unimodal system) or the ventral attention and limbic networks (heteromodal systems). The differential network composition across the fifteen overlapping nodes suggests a natural bipartition, with each half sitting towards either the lower or the higher end of the cortical hierarchy, respectively.

**Figure 4:**
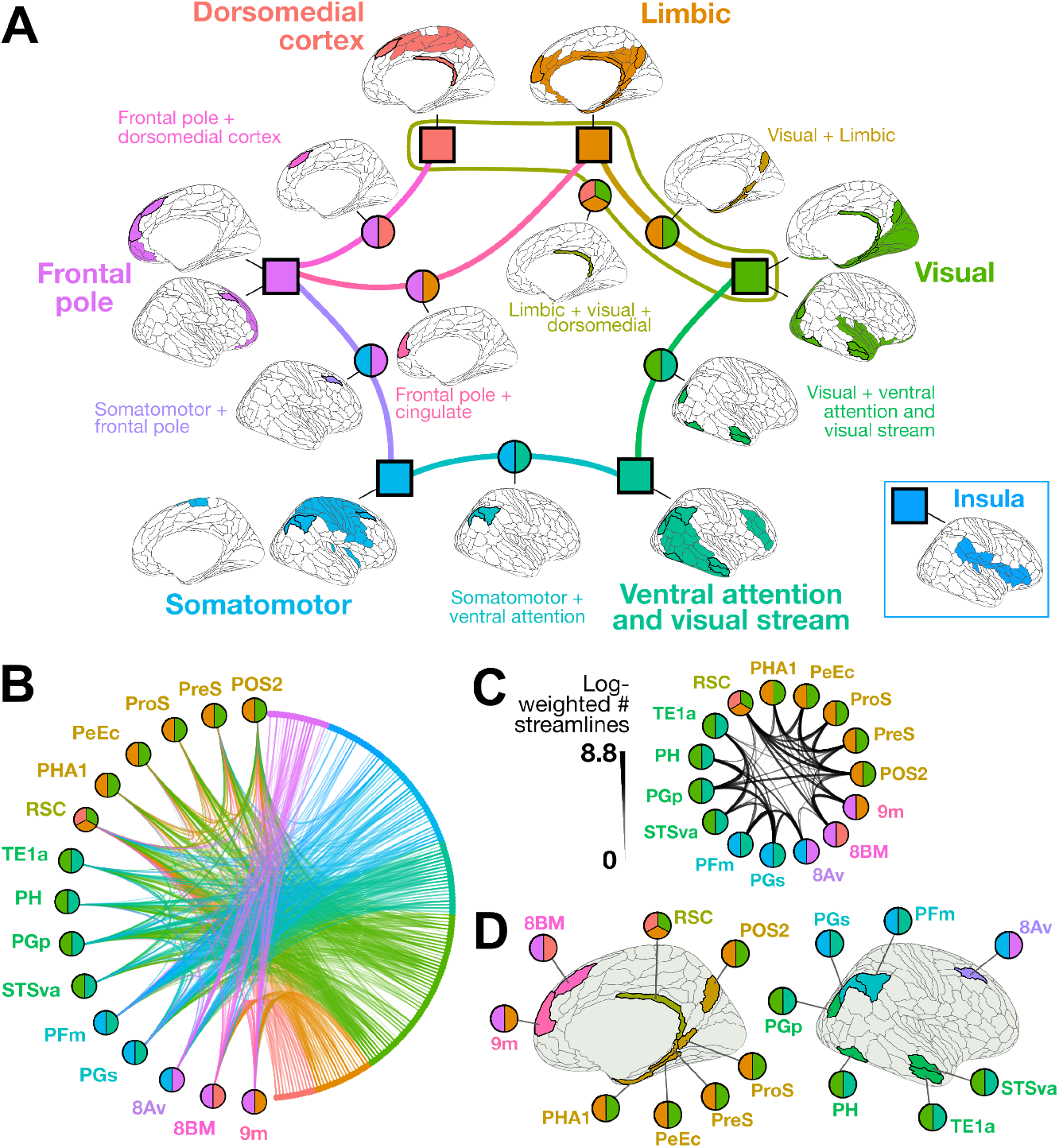
The overlapping community structure of the human structural connectome (right hemisphere) shows anatomical specificity and cross-network structural connectivity. **A** The `OSLOM_30` algorithm identified seven overlapping communities of the human structural connectome, which are represented with square nodes and corresponding brain maps with the 180-region parcellation from Glasser et al. [68]. Regions colored in a given map at each node were assigned to the corresponding module (and potentially to another module as well). The edges and their corresponding brain maps indicate regions with overlapping community assignments to two or more communities. Overall, the resulting communities are anatomically localized to the following regions: I. Dorsomedial cortex, II. Limbic, III. Visual, IV. Ventral attention + visual stream, V. Somatomotor, and VI. Insula, and VII. Frontal pole. Note that the ‘Insula’ module is positioned to the side as it does not share any overlapping connections to any of the other communities. **B** In this connectogram, every node is plotted on the edge of the circle, ordered by assigned structural module. Lines connecting nodes indicate connectivity between the pair of regions, with width and transparency showing the log-transformed streamline count. Fifteen brain regions were assigned to two different communities (‘overlapping regions’), distinguished on the left of the plot with two- or three-tone circles used to indicate the networks bridged by the corresponding region. The full structural connectome is summarized between these eleven overlapping regions (left) and the rest of the brain (right), with the latter colored by `OSLOM_30`-defined community as in **A**. Edges are colored according to the OSLOM community to which the connecting region on the right belongs, with edge weight and transparency mapping to the average number of tracts between the two regions. **C** The structural connectome between the fifteen overlapping regions is depicted, with line width and transparency indicating the log-transformed streamline count. **D** Each of the fifteen overlapping nodes is highlighted in color on the brain’s surface, annotated with the same multi-tone circles as in **A** to indicate shared community membership.

**Figure 5:**
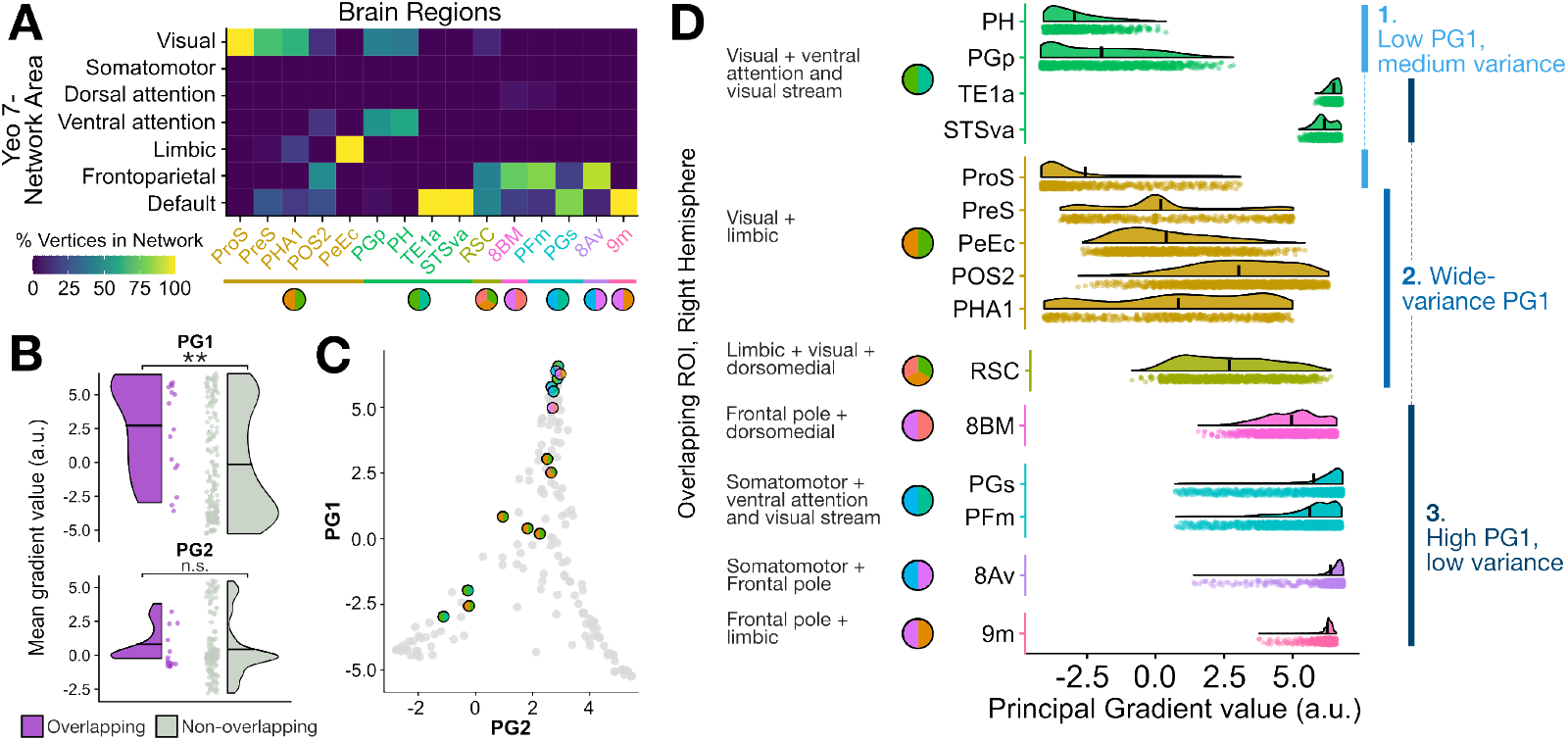
Regions participating in multiple structural communities generally sit higher in the human cortical hierarchy. **A**. For the fifteen cortical brain regions assigned to two or more modules, the percentage of vertices corresponding to each of the seven resting-state networks from [89] is shown as a heatmap. **B**. The mean first (PG1) and second (PG2) principal gradient values computed across all vertices per region are shown in raincloud plots for overlapping regions (purple, 15 regions) and non-overlapping regions (gray, 164 regions). ^**^, *P <* 0.01, Wilcox rank-sum test; n.s., *P >* 0.05. **C**. The mean PG1 versus PG2 values are plotted per brain region, with overlapping regions indicated by multi-tone circles as in **A. D**. For each group of overlapping nodes (e.g., ‘Visual + cingulate and parahippocampal’), the PG1 values are plotted for all vertices as raincloud plots. a.u., arbitrary units. In all plots, the black bar spanning each violin half represents the mean of the corresponding distribution. On the righthand side of raincloud plots, nodes have been grouped into one of three categories based on the patterns of PG1 value distributions.

We next quantitatively compared the location of each overlapping node along the cortical hierarchy by examining the first and second principal gradients (PGs) of functional connectivity [75] provided in the *neuromaps* atlas [76] (cf. Sec 2.6). As described in Margulies et al. [75], the first PG tracks a topographical hierarchy in the cortex, anchored by unimodal sensorimotor regions at the base and transmodal association regions at the top, with heteromodal regions supporting domain-general processing along the middle; whereas the second PG distinguishes between primary modalities by separating auditory and sensorimotor from visual regions. As shown in Fig. 5B, overlapping brain regions identified by `OSLOM_30` exhibit significantly higher first PG (PG1) values—averaged across all vertices per region—than non-overlapping counterparts (*P* = 0.004, Wilcox rank-sum test). This supports the notion that regions bridging two or more structural communities sit higher in the topographical hierarchy of the cortex on average. Although there was no statistically significant difference in PG2 values between overlapping versus non-overlapping nodes (*P* = 0.14, Wilcox rank-sum test), plotting nodes in PG1–PG2 space in Fig. 5C revealed that overlapping regions sit within the visual–transmodal axis, with a notable lack of regions in the auditory region (characterized by low PG1 and high PG2 values). This supports the notion that regions bridging two or more structural communities generally sit higher in the topographical hierarchy of the cortex.

The spread of PG1 values among overlapping regions is surprising, indicating that the nodes do not all sit narrowly at the top of the cortical hierarchy. Rather, the overlapping regions might be positioned across multiple hierarchical levels, which could allow them to act as intermediaries in integrating across systems that may sit at very different levels of the functional hierarchy. Indeed, Najafi et al. [17] found that overlapping regions exhibit greater functional diversity than those participating in only one network. While the overlapping regions did not exhibit more variance in vertex-wise PG1 values compared to non-overlapping regions (as shown in Fig. S4), we aimed to investigate the overall distribution of PG1 values across vertices in each overlapping region. As shown in Fig. 5D, we found interesting and distinctive patterns of vertex-wise PG1 values in overlapping regions, which we separated into three distinct patterns: (1) Low PG1 values with moderate variance, corresponding to *ProS, PH*, and *PGP*, which map primarily to the visual and ventral attention resting-state networks; (2) Moderate PG1 values with high variance, corresponding to heteromodal integration of domain-general processing, in the tri-modular *RSC* node and in four of the bi-modular ‘Visual + limbic’ nodes: *PreS, PeEc, POS2*, and *PHA1* ; and (3) High PG1 values with low to moderate variance, corresponding to transmodal association of higher-order functions, in the remaining nodes: *TE1a, STSva, 8BM, PGs, PFm, 8Av*, and *9m*. These distinctive patterns in the mean and variance of PG1 values suggest that while overlapping regions generally sit higher in the putative functional connectivity hierarchy, potentially enabling more synergistic systems-level processing—though this pattern is not universal across all overlapping regions. Rather, our findings suggest that the role of a given overlapping node within its complex network topology may depend upon the two or more communities it bridges, along with other structural and functional properties, from cytoarchitecture to neurotransmitter receptor densities [97, 98].

### 3.3 Comparing OSLOM with Louvain demonstrates the importance of allowing network-bridging overlaps

Having demonstrated that `OSLOM_30` generates biologically plausible structural networks bridged by overlapping nodes, we next asked how much additional information this overlapping modular network decomposition provides beyond that of a conventional, non-overlapping partition. In particular, we wanted to understand whether the overlapping nodes assigned by OSLOM exhibit distinctive nodal metrics of network integration relative to those of a non-overlapping partition, such as participation coefficient, *P*, and within-module strength (*z*-score), *z* [78]. As originally described in Guimerà and Nunes Amaral [78] and further extended to brain connectivity subsequently [99–101], comparing *P* and *z* can provide insight into the functional role that a given node may fulfill in its community, from peripheral hubs (low *z*, low *P*) through global connectors (high *z*, high *P*). Since the fifteen `OSLOM-30`-defined overlapping nodes exhibited many connections across communities and between each other (cf. Figs. 4B,C), we expected that they would sit in the ‘global connectors’ (high *z*, high *P*) space.

For comparison to the `OSLOM-30` overlapping partition, we selected the Louvain clustering method [77]— the most widely used community-detection method in network neuroscience [102]—to compute a non-overlapping modular decomposition, from which *z* and *P* were derived for each node (as described in Sec. 2.7). Using a representative decomposition with the resolution parameter *γ* = 1—which is the default in the Brain Connectivity Toolbox [81] and is supported by our robustness analyses in Fig. S5—we identified six non-overlapping Louvain communities, shown on the brain’s surface in Figs S5B,C. We also performed a robustness analysis with *γ* = 1.5 to enforce a seven-module (non-overlapping) decomposition, and the results were comparable to those obtained with *γ* = 1 (cf. Fig S5D-G), so we focus on the *γ* = 1 Louvain results hereafter. To characterize nodes according to Louvain versus OSLOM partitioning, we compared *P* versus *z* scores for each node. We show each node in *P*_Louvain_– *z*_Louvain_ space in Fig. 6B, where overlapping nodes (colored circles) are compared with non-overlapping nodes (gray circles). Most of the `OSLOM-30`-defined overlapping nodes exhibit *z*_Louvain_ *>* 0 (11 out of 15 nodes), indicating strong within-Louvain-module connectivity relative to other regions. However, only half of the overlapping nodes (8 out of 15) exhibit *P*_Louvain_ *>* 0.5, and even those do not stand out as exhibiting particularly high *P*_Louvain_ relative to non-overlapping nodes. For example, regions *8BM* and *9m*—judged as overlapping by `OSLOM-30`—exhibit strong within-Louvain-module connectivity (high *z*_Louvain_) yet sit towards the lower end of the *P*_Louvain_ distribution. This is surprising, as we expected regions that `OSLOM-30` identified as overlapping to show a greater spread of connectivity to nodes across multiple Louvain-defined modules (i.e., closer to ‘global connectors’, or at least ‘provincial hubs’).

**Figure 6:**
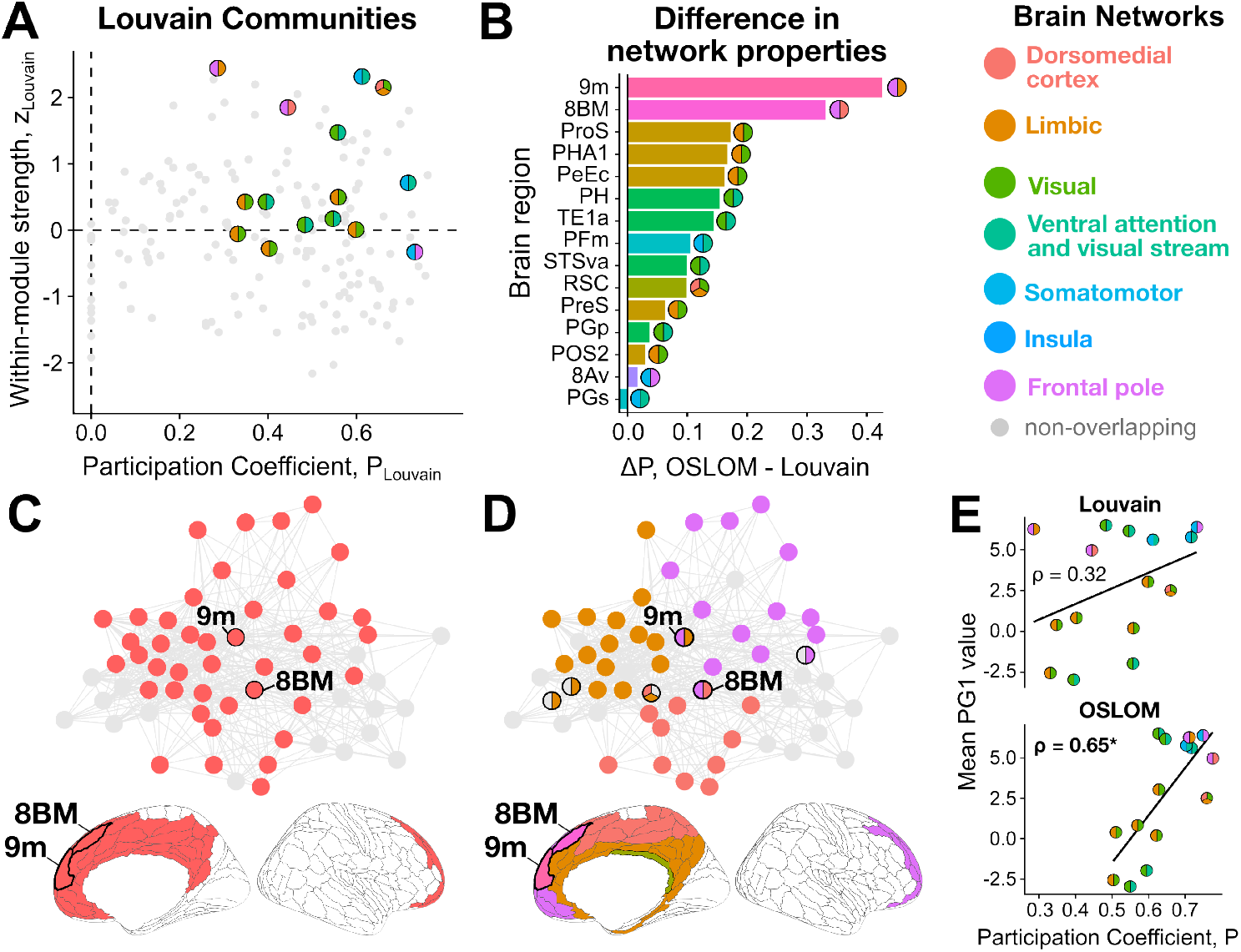
Overlapping nodes cannot straightforwardly be inferred from nodal metrics of a non-overlapping community decomposition. **A**. Scatter plots for *P* versus *z* for every node in the right hemisphere cortical connectome using the Louvain (non-overlapping) partition. The overlapping nodes (obtained by `OSLOM-30`) are marked in two-tone circles, with the two colors indicating the pair of communities bridged by the node in the OSLOM decomposition. **B**. The difference in *P*_OSLOM_ versus *P*_Louvain_ is compared for each OSLOM-identified overlapping region, ranked by the value of Δ*P*. **C**. Louvain assigns nodes *8BM* and *9m* to a community of 46 total nodes spanning frontal, cingulate, and dorsomedial cortices (n.b., only a subset of these nodes are shown that are structurally connected to *8BM* and/or *9m*), whereas **D**. OSLOM subdivides this set of nodes into three communities, collectively bridged by *8BM* and *9m*. The two-tone circles for *8BM* and *9m* indicate the two structural communities to which they were assigned. Circles with pink-gray or orange-gray coloring indicate overlap between the ‘Frontal pole’ or ‘Limbic and parahippocampal’ and other modules, respectively. Completely gray circles represent non-overlapping nodes. Nodes *8BM* and *9m* are annotated with bolded edges on the brain’s surface below the network plots in **C** and **D. E**. For each overlapping node, its mean PG1 value is compared with its participation coefficient from either Louvain or OSLOM partitioning. Spearman’s *ρ* is shown for each comparison; ^*^, *P <* 0.05.

To compare network properties with the `OSLOM-30` partition, we adapted the participation coefficient, *P*, to allow each overlapping node to be counted as part of its constituent two or three communities (noting that the specific module assignment label does not affect the value of *P*). With this adaptation (cf. Sec 2.7), *P* will increase by definition, so we focus on the relative differences in the magnitude of change (i.e., Δ*P*) in Fig. 6B. Regions *9m* and *8BM* stood out as the two regions with the greatest increase in *P* as measured by OSLOM relative to Louvain—prompting more detailed investigation of these regions. To investigate the structural changes in modular structure that may have underpinned these substantial increases in participation coefficient for 9m and 8BM, we further examined their ‘sub-network graphs’—comprised of the cortical regions sharing structural connections to one or both of *8BM* and *9m*. These subnetworks including *9m* and *8BM* are plotted in Figs 6C and D, for the Louvain and OSLOM modular decompositions, respectively. As shown in Fig. 6C for the Louvain decomposition, both *9m* and *8BM* were assigned to the same community, comprised in total of 46 nodes (of which 40 are included in this subnetwork) that collectively span frontal, cingulate, and dorsomedial cortices. We find that the same nodes depicted in the OSLOM decomposition in Fig. 6C are partitioned into two distinct communities that are bridged by *9m* (the ‘Frontal pole’ and ‘Limbic’ modules) and another two by *8BM* (the ‘Frontal pole’ and ‘dorsomedial cortex’).

These distinct modular decompositions demonstrate that OSLOM can disentangle multiple partially overlapping modules, capturing more complex yet subtle structure that is lost in a non-overlapping decomposition which cannot accommodate overlaps and therefore merges nodes into one larger module. While OSLOM detects different structures in the network, and thus yields different network nodal statistics like the participation coefficient, we next aimed to investigate whether the participation coefficient derived from Louvain or OSLOM may better capture other independently measured properties of brain organization among overlapping nodes, such position on the topographical cortical hierarchy [103]. As plotted in Fig. 6E for the fifteen overlapping nodes identified by OSLOM, the mean PG1 values (cf. Figs 5B-D) were significantly correlated with *P*_OSLOM_ (Spearman’s *ρ* = 0.65, *P* = 0.01) but not with *P*_Louvain_ (*ρ* = 0.32, *P* = 0.24). Similarly, we found no association between PG1 values and *P*_Louvain_ in our robustness analysis with *γ* = 1.5 (*ρ* = 0.37, *P* = 0.2; cf. Fig S5G). This indicates that the overlapping regions identified by OSLOM exhibit a greater degree of cross-module connectivity that positively correlates with position in the topographical hierarchy of the cortex. The same relationship holds when examining all 180 regions from the HCP–MMP1 right-hemisphere atlas [68]: as shown in Fig. S6, all mean PG1 values were significantly correlated with *P*_OSLOM_ (Spearman’s *ρ* = 0.33, *P* = 6 × 10^*−*6^) but not with *P*_Louvain_ (*ρ* = 0.01, *P* = 0.9). In other words, testing the hypothesis that the participation coefficient increases along the cortical functional hierarchy yields different conclusions using an overlapping versus non-overlapping decomposition. Only the OSLOM-derived participation coefficient *P* provides evidence for this hypothesis, with regions broadly increasing in their *P* from the unimodal end of the hierarchy up to the transmodal, associative end. Collectively, our comparison of metrics derived from OSLOM versus Louvain partitions underscores the utility of flexible overlapping decompositions for interpreting the resulting structural networks and derived nodal statistics, which are widely interpreted across the network neuroscience literature.

## 4 Discussion

This work introduces a general data-driven framework for selecting an overlapping community-detection algorithm (OCDA) for a given application. We further demonstrate how such overlapping modular decompositions can provide novel insights into the structure of the structure of human cortical connectivity. For a given target empirical dataset—here the structural connectome—the proposed method systematically compares how well each OCDA recapitulates the ground-truth overlapping community structure across benchmark networks generated from properties of the target network. This problem of calibrating methods to data relates to the concept of an algorithmic ‘footprint’ characterized in related work on other data types [104], that captures the differing regions (or ‘footprints’) in the data space in which different types of algorithms exhibit strong performance. Acknowledging that each OCDA makes different assumptions about network structure and inter-regional communication—and therefore that no OCDA can exhibit good performance on every type of network (cf. the ‘no free lunch’ theorem [36, 38])—our method circumvents the typical subjectivity in selecting an OCDA for a given problem, at the expense of constraints imposed on the simulated network ensemble by the benchmark generation process [47]. This demonstrates the importance of empirically tailoring OCDAs to a given network problem at hand, rather than relying on overly general statements about the relative performance of one algorithm over another [37]. We demonstrate an application of our approach to the structural connectome of the human cortex, where we select an appropriate OCDA algorithm (OSLOM) from an ensemble of brain-like networks. We then show how OSLOM quantifies new types of structure in brain networks, particularly by identifying overlapping regions that bridge across multiple biologically sensible modules and sit higher in the topographical cortical hierarchy. Further, the OSLOM decomposition yields nodal metrics that better distinguish these overlapping regions and align with independent measures of cortical organization, supporting the hypothesized hierarchical variation of nodal participation.

Using OSLOM to infer the overlapping community structure of the connectome, we obtain a useful and interpretable representation of the network structure with seven biologically plausible processing modules, collectively bridged by fifteen integrative overlapping nodes. All of the overlapping nodes sit at the spatial interface between their constituent and adjacent structural communities, suggesting they form connectivity ‘bridges’. Prior work suggests that higher-order integrative regions like *PFm* and *PFs*—which OSLOM identified here as spanning the ‘Somatomotor’ and ‘ventral attention and visual stream’ networks—serve as important functional and cytoarchitectonic transition zones between neighboring brain areas [105, 106]. In addition to *PFm* and *PFs*, regions *8BM* and *9m* in the dorsomedial prefrontal cortex also exhibited among the highest principal gradient of functional connectivity (PG1) values [75] of the regions we evaluated (cf. Fig. 5C). For these nodes, the structural connectivity overlaps may provide physical substrates for higher-order functional integration. Region *8BM*, which spans the ‘frontal pole’ and ‘dorsomedial cortex’ communities and sits in the frontoparietal resting-state network [89], has previously been highlighted as a higher-order cognitive control hub [107]. Moreover, Assem et al. [92] showed that *8BM, PFm*, and *PGs* are also components of the ‘multiple-demand’ network that orchestrates a variety of cognitive tasks, underlying flexible organization and cognitive control. Our findings are therefore consistent with the hypothesis that overlapping regions span multiple structural communities to enable functional integration.

However, some overlapping nodes’ PG1 values position them closer to heteromodal limbic regions (e.g., *POS2* and *RSC*) or even lower-order unimodal sensorimotor regions (e.g., *PH* and *PGp*). Despite sitting lower in the first putative topographical hierarchy of the cortex than other overlapping nodes, *POS2* (a component of the parieto-occipital sulcus in the HCP–MMP1 atlas [68]) participates in individual-specific functional network overlaps and is distinct from neighboring regions based on its cortical microstructure, functional connectivity, and task activation [34]. *RSC* (standing for ‘retrosplenial cortex’ in the HCP–MMP1 atlas [68]) is noteworthy as the only region identified to bridge across *three* structural networks here: ‘Limbic’, ‘visual’, and ‘dorsomedial cortex’. The RSC exhibits dense inter-network connectivity [108] and plays a critical integrative role in synthesizing viewpoints to process one’s environment [109, 110]. Taken together with the literature, our findings suggest that *POS2* and *RSC* serve as integrative transmodal hubs that bridge two or three distinct networks, respectively. For the lower-PG1 regions—*PH* in the temporo-occipital junction and *PGp* in the inferior parietal cortex—the potential role for structural overlap is less clear from the literature. *PGp* is a part of the angular gyrus (AG) believed to mediate semantic processing [111, 112], although evidence suggests it plays less of an ‘integrative’ role than other AG subregions [113]. Region *PH* contributes to visual processing from the foveal field [114], in line with the shared connectivity between the ‘visual’ and ‘ventral attention and visual stream’ networks identified herein.

Taken together, the differing patterns in functional connectivity principal gradient (PG1) values among the overlapping nodes suggest a wide spectrum of functions served by cross-network structural bridges, which likely depend on a variety of factors—including dynamic brain state, cytoarchitecture, receptor densities, and functional properties of the underlying brain systems. In support of this, Najafi et al. [17] described different ‘bridging’ behaviors exhibited across different overlapping regions based on the functional connectome, and Bijsterbosch et al. [34] found evidence for both inter-community coupling as well as ‘interdigitation’ (i.e., spatial interweaving of two networks) among areas of overlap between functional modules. Previous work has also suggested that such regions play a key role in information integration across brain modules and dynamical network states [1, 115–118]. Given this heterogeneity, future study is warranted to more fully characterize how the nature of overlapping regions relates to their own intrinsic structural, functional, and/or cytoarchitectonic properties, as well as those of the network communities they span.

Comparing intra- and inter-module network statistics, like *P* and within-module *z*, highlighted distinct network properties derived from an overlapping versus non-overlapping modular partition. Our results indicate that the inflexibility of the Louvain decomposition—which seeks the optimal nonoverlapping partition—can ‘force’ an overlapping modular structure into a single larger community in attempting to represent an underlying overlapping connectivity structure within the constraints of a non-overlapping partition, consistent with prior work with functional MRI [17]. In one example of this phenomenon (Fig. 6C), the Louvain partition consolidated regions *9m* and *8BM* into a singular 46-node community, with high *z* and low *P* (i.e., high intra-modular connectivity but low inter-modular participation). We also note that enforcing a seven-module Louvain decomposition with *γ* = 1 still placed *9m* and *8BM* in a singular, 32-node community. More generally, the fact that all the OSLOM-defined overlapping nodes were not distinctively concentrated in a specific area in the *z*_Louvain_ − *P*_Louvain_ space underscores how non-overlapping community-detection methods can yield suboptimal representations of the integrative, cross-module connectivity within a network structure. In other words, properties of a hard partition (like the within-module degree or participation coefficient in a non-overlapping decomposition) are insufficient to identify overlaps between communities. We note that *z* and *P* were both developed for non-overlapping community structure—thus requiring us to make slight modifications to these algorithms to account for overlapping nodes. Future work developing network characteristics that are readily amenable to both overlapping and non-overlapping algorithms would enable more rigorous direct comparison between two resulting decompositions.

Our methodology and results come with assumptions and limitations that should be interpreted in the context of this work. First, we assume that the generated ensemble of benchmark networks reflects the overlapping community structure of the target empirical network, and in turn that the top-performing OCDA on the benchmark ensemble will also perform well on the target network—such that ensemble properties are crucial to determining the ‘best’ OCDA. Moreover, we used the generative network model of Lancichinetti and Fortunato [47], which forces a power-law degree distribution that may not be optimally suited for many real-world networks—including the human connectome studied here. The model also does not incorporate other types of network properties, such as hierarchical community structure, which is a reported feature of brain-network organization [119]. As more sophisticated generative network models are developed that can incorporate a larger range of constraints, they can readily be incorporated into our framework to allow more precise calibration of OCDAs to target empirical data. In addition, given a generative model of overlapping community structure in networks, we had to make subjective choices about how to sample from the corresponding distribution over networks numerically. Here we sampled from sensible ranges of the relevant parameters—including community size distribution *τ*_2_, mixing parameters *µ*_*t*_ and *µ*_*w*_, and the number of overlapping nodes, *O*_*n*_—although these remain choices that can affect the assessment of the resulting OCDAs, and could be explored more thoroughly in future work.

Other limitations that should be considered with this work reflect inherent limitations with diffusion MRI-estimated structural connectomes. While there is empirical evidence that the probability for a connection between two regions exponentially decreases with the inter-region distance [86], the ‘distance bias’ in diffusion preprocessing can further decrease this probability—arising in part from the fact that reconstructing longer tracts introduces more steps for algorithmic errors [120]. Indeed, the seven distinct modules identified by OSLOM in the present study are generally spatially contiguous, with the exception of the ‘Visual’ and ‘Ventral attention and visual stream’ networks. Another potential limitation is that the average overlapping node occupies significantly more cortical surface area than the average non-overlapping node. A larger surface area can lead to an increased number of streamlines assigned to that region in the tractography estimation process [121, 122], and the overlapping regions do indeed have a higher degree on average, though we note that the overlapping nodes in this study were identified based on cross-module connectivity rather than the nodal degree. However, both region size and parcellation atlas can affect nodal statistics and the spatial location of overlapping regions within the brain [123, 124], so future work could expand to compare different parcellation schemes.

The method introduced here can be used to tailor overlapping community detection methods to network data, facilitating the additional insights that OCDAs can provide to diverse application areas. In our application to the human cortical connectome, our results highlight the importance of uncovering meaningful overlapping network partitions that better capture how the brain balances functional integration and segregation on a static backbone of structural connectivity. Future work could map out such a space for empirical networks with overlapping modular structure [125, 126] to more comprehensively characterize which parts of network space are best suited to which OCDAs. With this work, we also provide a comprehensive and extendable code base for overlapping community detection, which we hope will be expanded upon in future work, incorporating improvements in both new types of overlapping community-detection algorithms (including those that incorporate higherorder interactions [93]) and in overlapping benchmark network-generating algorithms.

## Acknowledgments

The authors thank Mac Shine, Isabella Orlando, Joshua Tan, and Christopher Whyte for their valuable input on data visualization and biological interpretation.

## Supplementary figures for ‘Tailoring overlapping community detection methods to the brain’s structural connectome’

**Figure S1:**
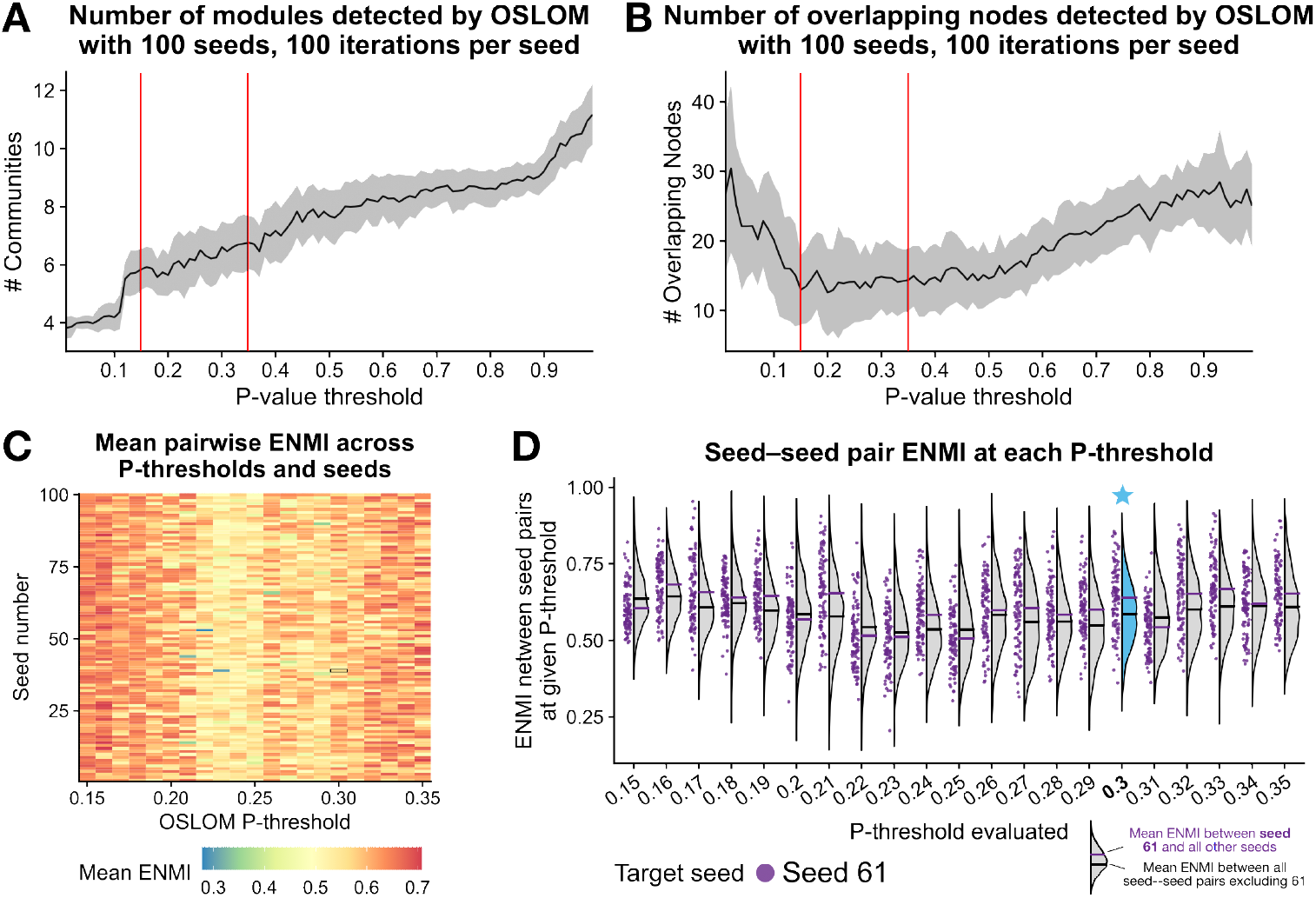
Parameter sweeps support the selection of *P* = 0.3 and seed number 61 for OSLOM. **A**. For each *P*-value threshold from 0.01 to 1, the mean number of communities that OSLOM identified from the empirical human cortical connectome is shown as a line plot. The shaded gray ribbon indices *±* 1 SD and the red lines highlight the 0.15 ≤ *P* ≤ 0.35 range. **B**. For each *P*-value threshold from 0.01 to 1, the mean number of nodes that OSLOM identified as overlapping across two or more communities is shown as a line plot. The shaded gray ribbon indices *±* 1 SD and the red lines highlight the 0.15 ≤ *P* ≤ 0.35 range. **C**. For the 0.15 ≤ *P* ≤ 0.35 range, the mean pairwise ENMI between each seed and all 99 other evaluated seeds is plotted as a heatmap. The color range indicates that the mean ENMI values range between [0.27, 0.71], with an average of 0.58 *±* 0.06 (arbitrary units). **D**. For the final seed selection (61), the pairwise ENMI between seed 61 and each of the 99 other seeds is depicted as a raincloud plot. Specifically, for each *P*-threshold from 0.15 ≤ *P* ≤ 0.35, there are 99 purple points corresponding to the ENMI between seed 61 and each of the 99 other seeds. The gray violins correspond to the ENMI distributions across all 4950 pairs of seeds from 1 to 100 (excluding self–pairs). Within each violin, the black bar measures the mean ENMI between all seed–seed pairs excluding seed 61, while the purple bar indicates the mean ENMI between seed 61 and all other seeds. The violin at *P* = 0.3 is highlighted in blue with a star as this is the final *P*-threshold choice in combination with seed 61, which yielded a mean ENMI of 0.64 *±* 0.11, compared to the average across all other seeds of 0.59 *±* 0.10.

**Figure S2:**
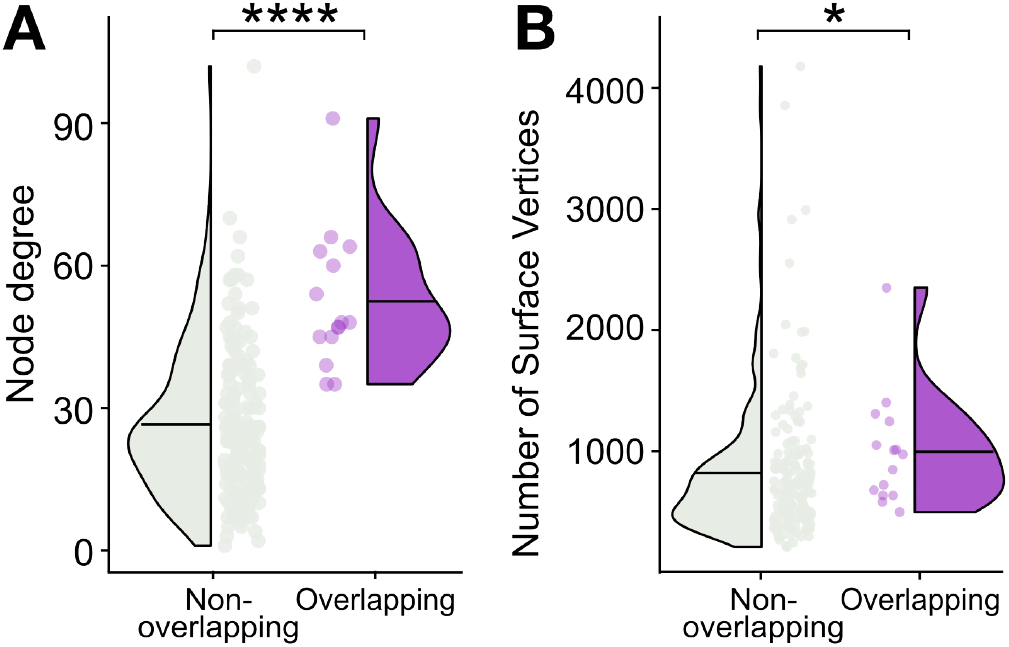
Overlapping regions identified by OSLOM_30 have a greater degree and are larger than non-overlapping regions on average. **A**. The distributions of nodal degree are shown for overlapping regions (purple, 15 regions) and non-overlapping regions (grey, 164 regions) as raincloud plots. ^****^, *P* = 3 × 10^*−*7^, unpaired Wilcox rank-sum test between overlapping vs. non-overlapping vertex counts. **B**. The distributions of cortical surface vertices contained within each overlapping region (purple, 15 regions) or non-overlapping region (grey, 164 regions) are shown as raincloud plots. ^*^, *P* = 0.03, unpaired Wilcox rank-sum test between overlapping vs. non-overlapping vertex counts.

**Figure S3:**
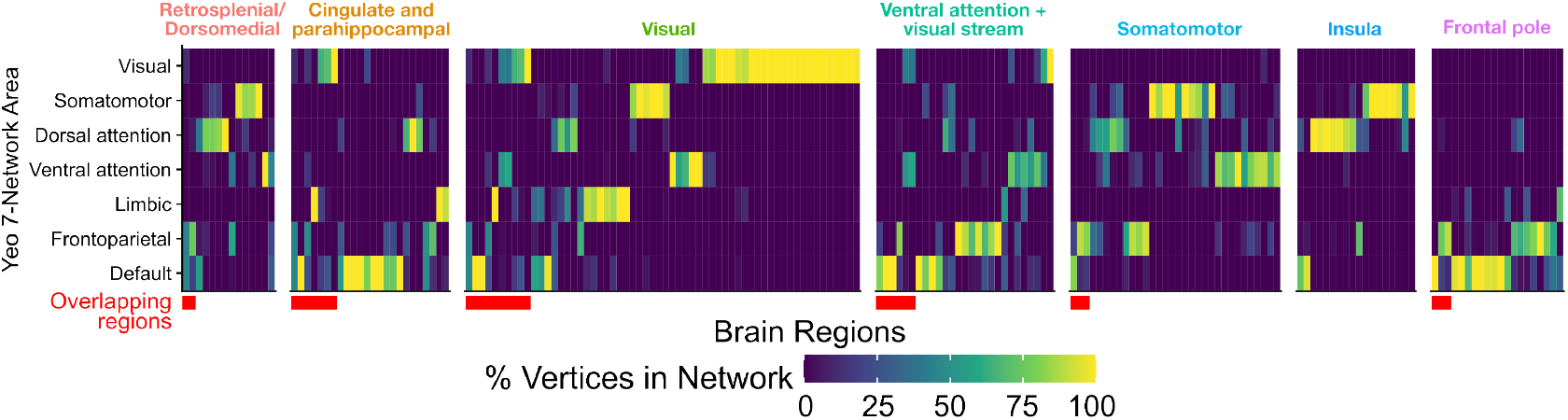
For each brain region, the proportion of vertices mapping to each of the 7-Network areas from Yeo et al. [89] was computed, with the percentages indicated as heatmap tile values. Brain regions are grouped along the *x*-axis by OSLOM-identified structural module, with the ‘overlapping’ regions to the left of each cluster (annotated by the red bar below the heatmap). Note that overlapping regions were included as columns in each of the corresponding OSLOM modules, such that each overlapping region appears two or three times in the heatmap.

**Figure S4:**
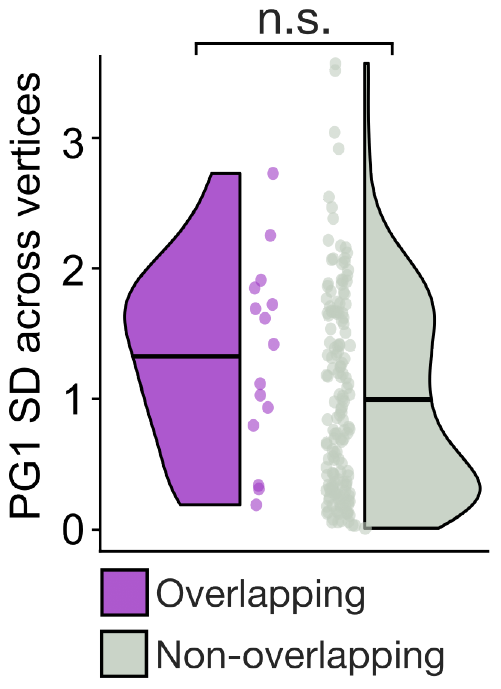
There is no difference in the vertex-wise SD of first principal gradient (PG1) values between overlapping vs. non-overlapping regions. The distributions of PG1 SD (across vertices) within each region are shown as raincloud plots for overlapping regions (purple, 15 regions) or non-overlapping regions (grey, 164 regions). n.s., *P >* 0.05, unpaired Wilcox rank-sum test between overlapping vs. non-overlapping PG1 SD values.

**Figure S5:**
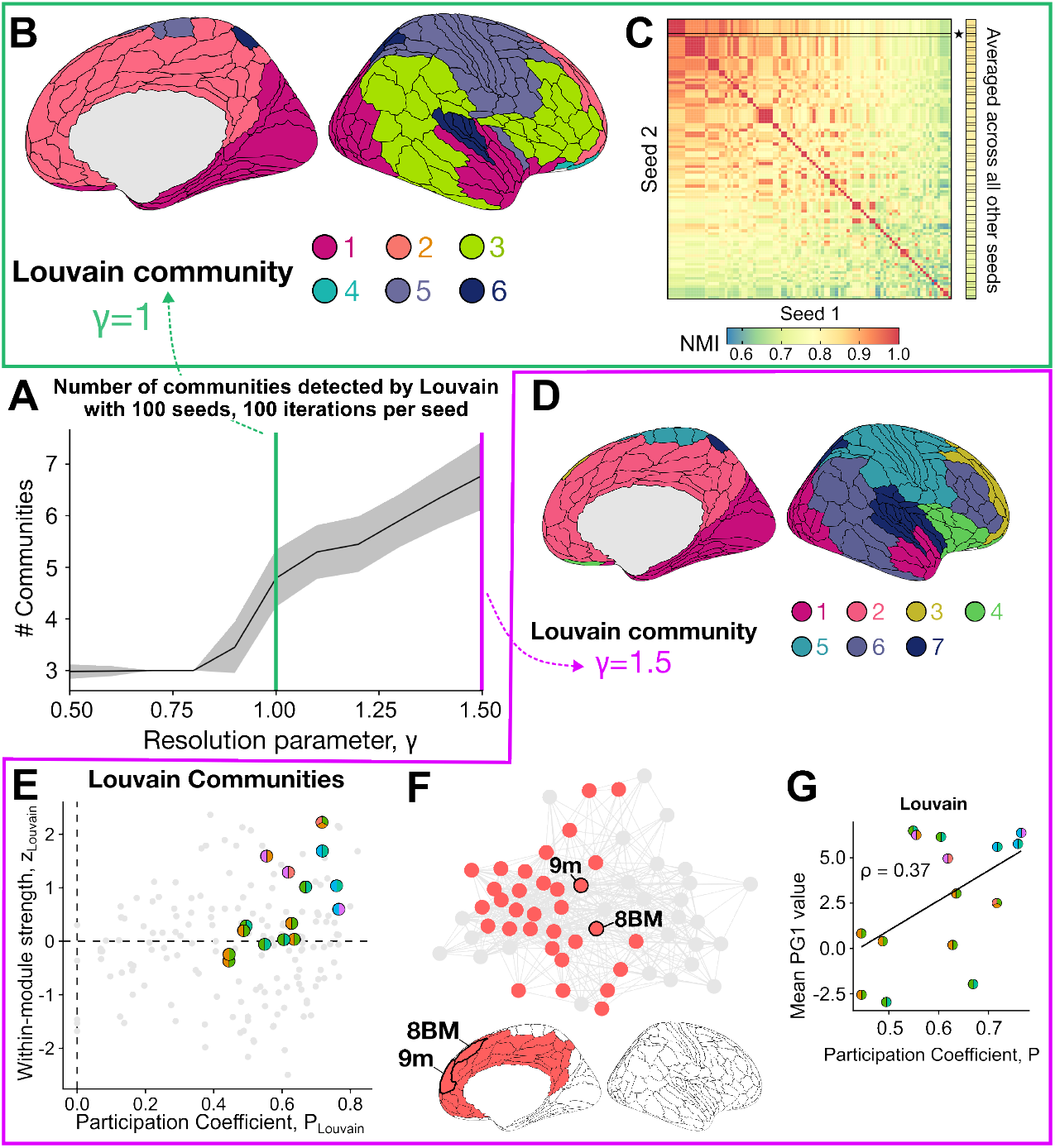
Louvain partitioning with *γ* = 1 yields a representative six-community decomposition, with comparable properties to that of a seven-community decomposition with *γ* = 1.5. **A**. For the resolution parameters *γ* = [0.5, 0.6, … 1.5], Louvain partitioning was performed using 100 initialization seeds each. The mean number of detected communities per *γ* value is plotted as a black line, with the shaded ribbon indicating ±1 standard deviation. The green line at *γ* = 1 is included to guide visual interpretation for this threshold selection parameter, with a representative six-community decomposition projected onto the cortical surface in **B. C**. The result in **B** corresponds to the seed number 98, which was selected based on its maximal normalized mutual information (NMI) value to all other evaluated seeds, as shown in the heatmap. The pairwise NMI values are depicted as a heatmap, with the selected seed (98) outlined in bold with a star. **D**. To compare our seven-community OSLOM decomposition with a seven-community Louvain decomposition specifically, we also evaluated a maximally representative seed (seed=91) with the threshold *γ* = 1.5. These seven communities are projected onto the right cortical surface here. **E**. Scatter plots for *P*_Louvain_ versus *z*_Louvain_ for every node in the right hemisphere cortical connectome, using *γ* = 1.5. The overlapping nodes (obtained by `OSLOM-30`) are marked in two-tone circles, with the two colors indicating the pair of communities bridged by the node. **F**. Louvain (with *γ* = 1.5) assigns nodes *8BM* and *9m* to a community of 32 total nodes spanning frontal, cingulate, and retrosplenial/dorsomedial cortices (n.b., only a subset of these nodes are shown that are structurally connected to *8BM* and/or *9m*). Gray circles in the network plot represent nodes that are structurally connected to *8BM* and *9m* but not assigned to the same Louvain module. **G**. For each overlapping node identified by `OSLOM-30`, its mean PG1 value is compared with its *P*_Louvain_. Spearman’s *ρ* = 0.37, *P* = 0.17.

**Figure S6:**
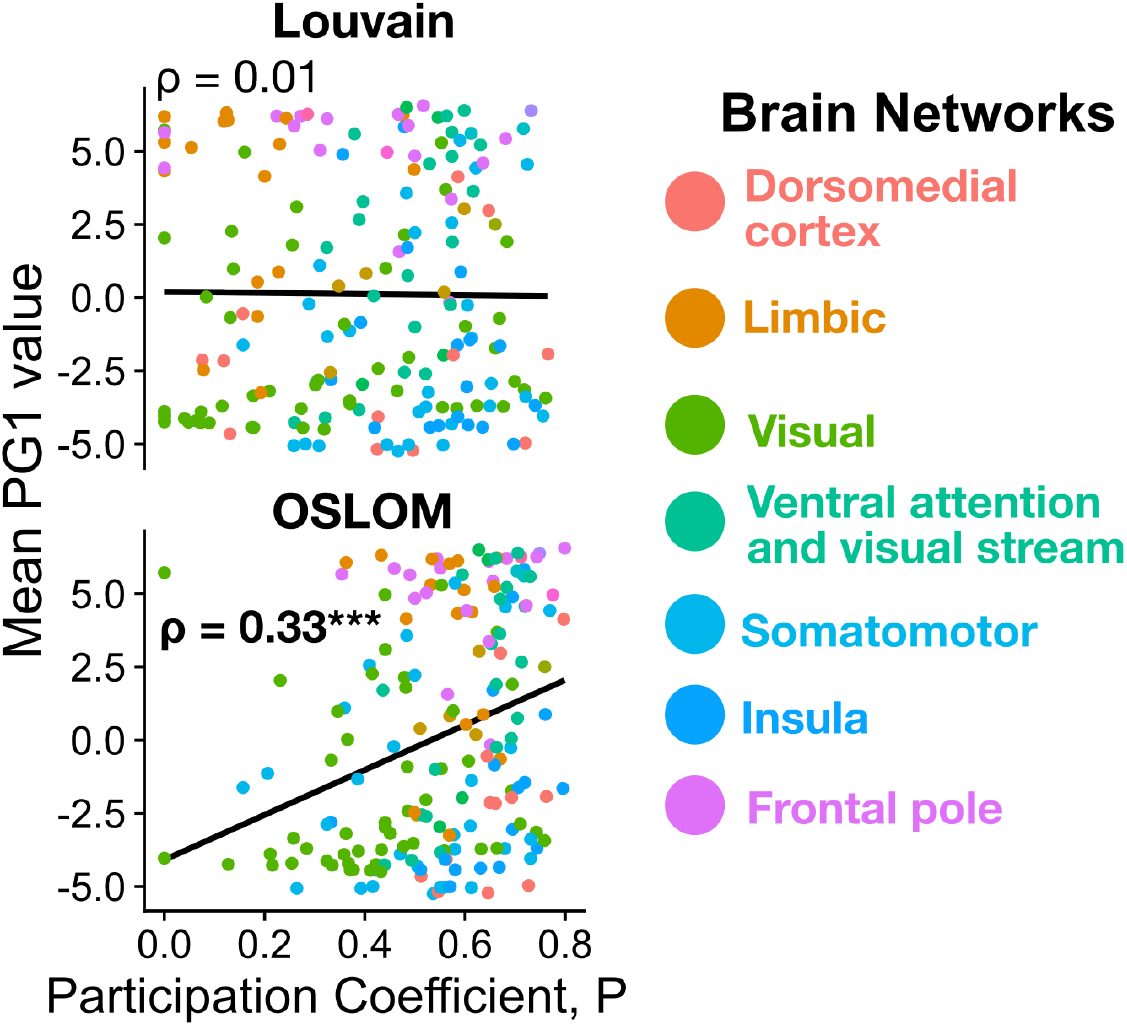
The participation coefficient from the OSLOM decomposition, but not the Louvain decomposition, is significantly correlated with the first principal gradient of functional connectivity (PG1) across the right hemisphere cortex. For each node in the 180-region HCP–MMP1 atlas, its mean PG1 value is compared with its participation coefficient from either Louvain (*P*_Louvain_) or OSLOM (*P*_OSLOM_) partitioning. Spearman’s *ρ* is shown for each comparison; ^***^, *P <* 0.001.

## Footnotes

1 https://github.com/NeuralSystemsAndSignals/OverlappingCommunityDetection

2 https://github.com/NeuralSystemsAndSignals/OverlappingCommunityDetection_HCP

